# HSP90 inhibitor NVP-HSP990 alleviates rotavirus infection

**DOI:** 10.1101/2023.06.15.545197

**Authors:** Yi Cao, Qingmin Zhu, Xiaoping Wu, Zhunyi Xie, Chengying Yang, Yanyan Guo, Dongwei Meng, Xinyuan Zhou, Yuzhang Wu, Jintao Li, Haiyang He

## Abstract

Rotavirus (RV) infection is a significant cause of hospitalization and mortality in infants and young children. Although conventional symptomatic treatments usually appear effective, tens of thousands of infants and young children still die each year due to the absence of safe and effective anti-RV drugs. Heat shock protein 90 (HSP90) is usually required for efficient viral infection; however, due to unsatisfactory antiviral efficacy and toxicity, there has been no HSP90-targeting agents applied for clinical antiviral therapy currently. Here, we demonstrated that NVP-HSP990, a novel small-molecule HSP90 inhibitor with excellent oral bioavailability and brain penetration, was a potent inhibitor of RV infection with much bigger selectivity index (SI) than traditional HSP990 inhibitors. NVP-HSP990 potently inhibited RV replication *in vitro* without blocking infection establishment. NVP-HSP990 remarkably restored gene expressions of most KEGG pathways disturbed by RV infection in intestinal cells, except some inflammatory pathways (IL-17, TNF, etc.). To be noted, NVP-HSP990 significantly altered gene expressions in MAPK signaling pathway and inhibited RV-induced activation of MAPK as well as disruption of tight junctions in Caco-2 cells. More importantly, NVP-HSP990 effectively alleviated RV diarrhea, competently inhibited RV replication, and obviously prevented pathological lesions of intestine in BALB/c suckling mice. Therefore, our results suggested that NVP-HSP990 can be a promising antiviral drug candidate against RV infection.

## 1. Introduction

Rotavirus (RV) infection is the leading cause of diarrhea in infants and young children(1). According to the latest global survey by WHO, RV causes approximately 0.64% of hospitalizations and around 208,009 deaths of children under 5 years of age in 28 low- and middle-income countries(2). RV is a double-stranded RNA virus that belongs to the Reoviridae family. It infects enterocytes and causes dysfunction of enterocytes and disorder of the enteric nervous system, which leads to severe dehydrating diarrhea and vomiting. RV infection also attacks other important organs, particularly the brain, leading to epileptic seizures and other central nervous system disorders(3–5). In clinic, symptomatic treatments and gastrointestinal protective agents are usually the main choices for RV infection, while conventional antiviral drugs like nucleoside analogues and type I interferon are not typically used to treat RV infection due to their adverse side-effects(6, 7), which sometimes leads to uncontrolled viral infection and even mortality. Therefore, there is an urgent need to research safe and effective antiviral drugs for RV infection.

Heat shock protein 90 (HSP90) is a chaperone protein found in both eukaryotes and bacteria. It has four isoforms (α, β, GRP94, and TRAP1) that are located in different parts of cells(8, 9). HSP90 promotes viral replication by regulating cell signaling systems or by direct interacting with viral proteins such as Hepatitis B virus reverse transcriptase, Hepatitis C virus non-structural protein 5A, and Influenza virus A RNA-dependent RNA polymerase(10–13). HSP90 is involved in the entry of RV into certain tumor cell lines(14–16), and contributes to RV replication *in vitro*(17, 18). Therefore, it seems that HSP90 inhibitors would be promising broad□spectrum antiviral drug candidates(19); nevertheless, in contrast to the widespread application in anti-tumor therapies(20), no HSP90 inhibitors are in clinical use for antiviral therapies currently, probably due to unsatisfactory antiviral efficacy and undesirable toxicity.

NVP-HSP990 is a novel small-molecule HSP90 inhibitor with a distinct structure from the natural HSP90 inhibitor Geldanamycin (GA) and its derivative Tanespimycin (17-allylamino-17-demethoxygeldanamycin, 17-AAG)(21–23). NVP-HSP990 exhibits high oral bioavailability and binds to the N-terminus of both HSP90α and HSP90β with high affinity(24); in addition, NVP-HSP990 is able to penetrate the blood-brain barrier and very low dose of NVP-HSP990 exerts potent antiepileptic activity(25, 26). These advantages make NVP-HSP990 a potential antiviral drug candidate, especially for viruses that offense the brain. However, currently there is limited experimental data on the effects of NVP-HSP990 on viral infection. In this study, we evaluated the inhibitory effects of NVP-HSP990 on RV infection, and our results suggested that NVP-HSP990 is a promising potential antiviral agent for RV infection.

## 2. Materials and Methods

### Cell culture, viral infection, and drug administration

MA104 cells were provided by Dr. Elschner (Friedrich-Loeffler-Institute). Caco-2 and HT-29 cells were from ATCC and kept in our institute. All the cells were cultured in Dulbecco’s Modified Eagle Medium (DMEM) (Invitrogen) plus 10% (v/v) fetal bovine serum (FBS) (Gibco) at 37°C with 5% CO_2_. RV Wa and SA11 strains were from the China Center for Disease Control and Prevention (Beijing, China) and propagated in MA104 cells. For RV infection, the viruses were diluted in DMEM, activated with 10 μg/ml trypsin (Amersco) for 30 min at 37°C, and then added to target cells previously washed once with DMEM. After 1 h of incubation at 37°C, the inoculum was removed; the cells were then washed twice with DMEM and incubated with DMEM at 37°C till use. HSP90 inhibitors NVP-HSP990, GA, and 17-AAG (Selleck) were dissolved in DMSO and either administrated with indicated concentrations *in vitro* or orally administrated with indicated quantity plus 90% (vol/vol) corn oil.

### Virus preparation

RV-infected MA104 cells and the culture medium were frozen/thawed twice and centrifuged at 10,000 g for 20 min at 4°C for debris clearance. Supernatant of Wa strain was concentrated by centrifugation at 100,000 g with a SW28 rotor for 2 h at 4°C. For concentration of SA11 strain, NaCl was added to the supernatant of SA11 strain to a final concentration of 0.5 M, then Polyethylene glycol (PEG, MW6,000, Sangon Biotech) was added to a final concentration of 8% (w/v) with stirring overnight at 4°C, followed with centrifugation at 10,000 g for 2 h at 4°C. Both the virus pellets were suspended in TNC buffer (50 mmol/L Tris, 140 mmol/L NaCl, 10 mmol/L CaCl_2_, pH 8.0), filtered with 0.22 μm filters and subjected to sucrose density gradient centrifugation. Viruses were titrated and stored in aliquots at –70°C.

### Cytotoxicity assay

To evaluate the cytotoxicity of different HSP90 inhibitors *in vitro*, 10000 MA104, Caco-2 or HT-29 cells were seeded in 96-well plates and cultured for 12 h, then the indicated concentrations of NVP-HSP990, GA, or 17-AAG were added into the culture medium respectively, while a cell-free control was also set. 10 μl CCK-8 reagents (Beyotime) were added into each well 24 h after drug administration, and OD450 absorbance of each well was detected 3 h later using a Microplate Reader (Gene Company Limited). Relative cell viability rate = OD450 (drug-treated cells - cell free control) / OD450 (0 μM drug-treated cells - cell free control). The 50% cytotoxic concentration (CC50) was calculated by the GraphPad Prism 9.0 software.

### Plaque formation assay (PFA)

PFA was carried out as described(27) with some modification. Briefly, confluent monolayers of MA104 cells in 24-well tissue culture plates were infected with 100 μl supernatants containing infectious RV particles which were serially diluted (10^1^, 10^2^, 10^3^, and 10^4^) with DMEM containing 1 μg/ml trypsin and penicillin (10,000 U/ml)-streptomycin (5 μg/ml) (DMEM-T) for 1 h at 37°C. Then 400 μl agarose overlay containing DMEM-T was added for 5 days. The cells were then fixed with 4% paraformaldehyde, followed by staining with 1% crystal violet for counting of plaque forming units.

### In vitro assay of viral inhibitory effects of drugs

For analysis of effects of different HSP990 inhibitors on RV reproduction *in vitro*, MA104, Caco-2, and HT-29 cells were infected with RV Wa or SA11 strains (MOI=1) and treated with indicated serial concentrations of NVP-HSP990, GA, or17-AAG. At 24 h p.i., the infected cells and culture medium was collected, frozen/thawed twice and subjected to centrifugation at 1,000 g for 3 min. Then, the supernatant was subjected to analysis of viral load by PFA, and the 50% inhibitory concentration (IC50) of virus reproduction was calculated with GraphPad Prism 9.0 software.

### RNA-sequencing (RNA-seq) analysis

Caco-2 cells growing on 6-well culture plate were mock infected with PBS or infected with RV Wa or SA11 strains (MOI=3), then the cells were cultivated with DMEM containing 100 nM NVP-HSP990 or equal volume of DMSO as control. At 24 h p.i., the Caco-2 cells were harvested and lysed in Trizol (Life Technologies) for RNA extraction. Total RNA was extracted using Trizol reagent kit (Invitrogen, Carlsbad, CA, USA) according to the manufacturer’s protocol. RNA quality was assessed on an Agilent 2100 Bioanalyzer (Agilent Technologies, Palo Alto, CA, USA) and checked using RNase free agarose gel electrophoresis. After total RNA was extracted, eukaryotic mRNA was enriched by Oligo(dT) beads. Then the enriched mRNA was fragmented into short fragments using fragmentation buffer and reversely transcribed into cDNA by using NEBNext Ultra RNA Library Prep Kit for Illumina (NEB #7530, New England Biolabs, Ipswich, MA, USA). The purified double-stranded cDNA fragments were end repaired, A base added, and ligated to Illumina sequencing adapters. The ligation reaction was purified with the AMPure XP Beads(1.0X).And polymerase chain reaction (PCR) amplified. The resulting cDNA library was sequenced using Illumina Novaseq6000 by Gene Denovo Biotechnology Co. (Guangzhou, China). Raw data (raw reads) of fastq format were further filtered by fastp (version 0.18.0). In this step, clean data (clean reads) were obtained by removing reads containing adapter, reads containing ploy-N and low-quality reads from raw data. Short reads alignment tool Bowtie2 (version 2.2.8) was used for mapping reads to ribosome RNA (rRNA) database. The rRNA mapped reads then will be removed. The remaining clean reads were further used in assembly and gene abundance. Reference genome Homo sapiens Ensembl_release110 and gene model annotation files were downloaded from genome website directly. Index of the reference genome was built using HISAT2.2.4 and paired-end clean reads were aligned to the reference genome using HISAT2.2.4. The mapped reads of each sample were assembled by using StringTie v1.3.1 in a reference-based approach. For each transcription region, a FPKM (fragment per kilobase of transcript per million mapped reads) value was calculated to quantify its expression abundance and variations, using RSEM software. Differential expression analysis was performed using the DESeq2 R package (1.16.1), RNAs differential expression analysis was performed by DESeq2 software between two different groups. The genes/transcripts with the parameter of false discovery rate (FDR) below 0.05 and absolute fold change≥2 were considered differentially expressed genes/transcript.

### Flow cytometry analysis

Differentially-treated MA104 or Caco-2 cells were infected with RV Wa strain (MOI=1). At 20 h p.i. the infected cells were digested with 0.25% trypsin-EDTA, the cells were fixed with 4% paraformaldehyde for 15 min, permeabilized with 1% Triton-X-100 for 7 min, quenched with 50 mM NH_4_Cl for 10 min, and washed for 3 min in 0.01 M PBS twice. Cells were stained with FITC-labeled goat anti-RV polyclonal antibodies (0503, Virostat) (1:100) for 1 h at room temperature, and the frequency of infected cells was analysed with BD Canton II flow cytometry (BD).

### Immunofluorescence assay (IFA)

Caco-2 cells growing on coverslips were mock infected with PBS or infected with RV Wa or SA11 strains (MOI=1), then the cells were cultivated with DMEM containing 100 nM NVP-HSP990 or equal volume of DMSO (CTRL) for another 18 hours after infection. The cells were then washed with 0.01 M PBS, fixed in 4% paraformaldehyde for 20 min, followed by permeabilization in 0.5% Triton-X-100 for 10 min and blocking with 5% BSA for 1 h at room temperature. The cells were then incubated with rabbit anti ZO-1 monoclonal antibody (13663S, CST) (1:100) for 2 h at room temperature, washed twice with 0.01 M PBS, incubated with Cy3-conjugated goat anti rabbit antibodies (A0516, Beyotime) (1:500) and FITC conjugated goat anti RV antibodies (1:100) for 1 h at room temperature, and followed by 2-(4-Amidinophenyl)-6-indolecarbamidine dihydrochloride (DAPI) (5 μg/ml) staining for 10 min. The triple-stained cells were washed twice with 0.01M PBS and we captured images at using Zeiss Axio Imager A2.

### qPCR analysis

RV-infected Caco-2 cells or intestines form RV-infected suckling mice were lysed in Trizol (Life Technologies) for RNA extraction and RNA was further purified using RNeasy Plus Micro or Mini Kit (Qiagen). RNA was reverse transcribed Using HiScript® II Q RT SuperMix for qPCR (+gDNA wiper) Kit (Vazyme), and qPCR was performed using AceQ qPCR SYBR Green Master Mix (Vazyme) with corresponding primers on a CFX96 Touch Real-Time System (Bio-Rad). Primers used for amplifications are listed in Supplemental Table 1, from which human or mouse β-actin was selected as internal reference.

### Western blot

MA104 and Caco-2 cells were infected with RV (MOI=3) or mock infected with PBS at 80% confluence and treated with 100 nM HSP990 (+) or equal volume of DMSO as control (-) for 20 h. The cells were lysed with RIPA Lysis Buffer (Beyotime) plus protease inhibitor cocktail (Thermo). The cell lysates were collected in 1.5 ml EP tubes, sonicated 5 times for 15 s at 80 watts on ice, clarified by centrifugation at 12,000 g 10 min at 4°C, subjected to precast 4-20% gels for SDS-PAGE and then transferred to 0.22 μm PVDF membranes (Millipore). After blocking with 5% BSA, the membranes were incubated with rabbit mAbs to MAPK components (JNK (9252T, CST), Phospho-JNK (4668T, CST), p38 (8690T, CST), Phospho-p38 (4511T, CST), ERK1/2 (4695T, CST), Phospho-ERK (4370T, CST)), goat pAbs to RV particles (0501, Virostat), or with mouse mAb to β-actin (GB15001, Servicebio) as internal reference. Afterwards, the membranes were incubated with horseradish peroxidase (HRP)-conjugated anti-rabbit (7074S, CST), anti-goat (A0181, Beyotime), or anti-mouse (7076S, CST) IgG. Immunoreactive bands were visualized by using enhanced chemiluminescence’s substrate (BeyoECL Plus, Beyotime).

### RV diarrhea of suckling mouse model

For induction of RV diarrhea of suckling mice, 1×10^6^ PFU RV (SA11 strain) was orally inoculated to 7-days-old BALB/c suckling mice regardless of their sexes, as sex of the BALB/c suckling mice has no significant influence on RV diarrhea. We used a scoring system to evaluate fecal consistency and color as described(28): no stool (0 point); brown formed stool (1 point); brown soft stool (2 points); yellow soft stool (3 points); and yellow watery stool (4 points). Mice with scores of ≥ 2 meant diarrhea occurrence, while mice with scores of < 2 meant no diarrhea. Diarrhea scores of the infected suckling mice were examined every 24 h by examination of fecal material. All work based on this animal model was done according to the requirements of Third Military Medical University Animal Ethics Committee.

### In vivo RV inhibition of NVP-HSP990

7-days-old BALB/c suckling mice were orally inoculated with 1×10^6^ PFU RV SA11 strain. At 2 h p.i., the mice were treated with indicated doses of NVP-HSP990, GA, or 17-AAG, or with equal volume of DMSO as control. The diarrhea scores were recorded at indicated time points. The EC50 (concentration for 50% of maximal effect) of NVP-HSP990 on inhibition of RV diarrhea occurrence or on reduction of diarrhea scores was calculated using the GraphPad Prism 9.0 software.

To test the effect of NVP-HSP990 on RV replications *in vivo*, 7-days-old BALB/c suckling mice were orally inoculated with 1×10^6^ PFU RV SA11 strain or PBS as mock infection. At 2 h p.i., these suckling mice were treated with 1 mg/Kg NVP-HSP990 or equal volume of DMSO as control, and then were sacrificed by CO_2_ asphyxiation at 16 h p.i.. Small intestines (including Duodenum, Jejunum, and Ileum) of these mice were homogenated in 0.3 ml DMEM, followed by centrifugation at 12,000 g for 5 min. The supernatant was collected for virus titration with PFA or for assay of viral antigens with Enzyme-linked immunosorbent assay (ELISA) using goat anti RV pAbs (0501 and 0503, Virostat) and RV antigen ELISA kit (CUSABIO) following the instructions. For qPCR analysis of VP6 expression, the intestines were longitudinally split and washed in 1 ml DMEM each by shaking for 30 s. Then, the tissues were cut into 0.5 cm segments, incubated in Hank’s buffer containing 2.5 mM EDTANa_2_ and 1 mM DTT at 37°C with rotation at 200 rpm for 30 min, filtered with 70 μm strainer. Then the epithelial cells were pelleted by centrifugation at 500 g for 5 min, purified with 20% Percoll, and lysed in Trizol for RNA extraction and further qPCR analysis.

### Histopathology

7-days-old BALB/c suckling mice were orally inoculated with 1×10^6^ PFU RV SA11 strain or PBS as mock infection. At 2 h p.i., these suckling mice were treated with 1 mg/Kg NVP-HSP990 or equal volume of DMSO as control, and then were sacrificed by CO_2_ asphyxiation at 16 h p.i.. The ileum of the suckling mice were fixed in 4% paraformaldehyde for 24 h, washed with PBS twice, and followed by paraffin embedding and slicing at 5 μm thickness. The sections were stained with hematoxylin and eosin (Beyotime, China) and observed with an Olympus BX51 microscope.

### Statistical analysis

Data were presented as means ± s.e.m. Statistical analysis was performed with Prism 9.0 (GraphPad). One-way analysis of variance (ANOVA) test was used for comparisons between multiple groups. Two-way ANOVA test was used for comparisons of grouped data. *P* values < 0.05 were considered significant (\**P* < 0.05; \*\**P* < 0.01; \*\*\**P* < 0.001; \*\*\*\**P* < 0.0001); ns: not significant, *P* > 0.05.

## 3. Results

### 3.1 Antiviral activity of NVP-HSP990 in vitro

NVP-HSP990 is structurally different from the traditional HSP90 inhibitors GA and 17-AAG and is much smaller (Supplementary Fig. 1). Using CCK-8 analysis, we compared the cytotoxicity of these HSP990 inhibitors on human intestinal epithelial cell line Caco-2, rhesus monkey embryo kidney cell line MA104, and human intestinal epithelial cell line HT-29, which are all susceptible to RV infection. NVP-HSP990 showed much less cytotoxicity than GA and 17-AAG, with CC50 (concentration of cytotoxicity 50%) values of 64.05 μM to Caco-2 cells, 62.25 μM to MA104 cells, and 59.89 nM to HT-29 cells (Fig. 1A). To test the anti-RV effect of NVP-HSP990 *in vitro*, Caco-2, MA104, and HT-29 cells were infected with human RV Wa strain or simian RV SA11 strain with a multiplicity of infection (MOI) of 1 and cultured in DMEM containing a series of concentrations of HSP990 inhibitors for 24 h. It was shown that NVP-HSP990 inhibited RV reproduction with IC50 (half maximal inhibitory concentration) values of 4.046 nM (Wa strain) and 4.686 nM (SA11 strain) in Caco-2 cells, 1.067 nM (Wa strain) and 0.912 nM (SA11 strain) in MA104 cells, and 6.620 nM (Wa strain) and 3.964 nM (SA11 strain) in HT-29 cells, which were all far smaller than those of GA and 17-AAG (Fig. 1B). Therefore, the selectivity index (SI) of NVP-HSP990 was much smaller than which of GA or 17-AAG in all tested cells in RV infection (Fig. 1C).

**Fig. 1.**
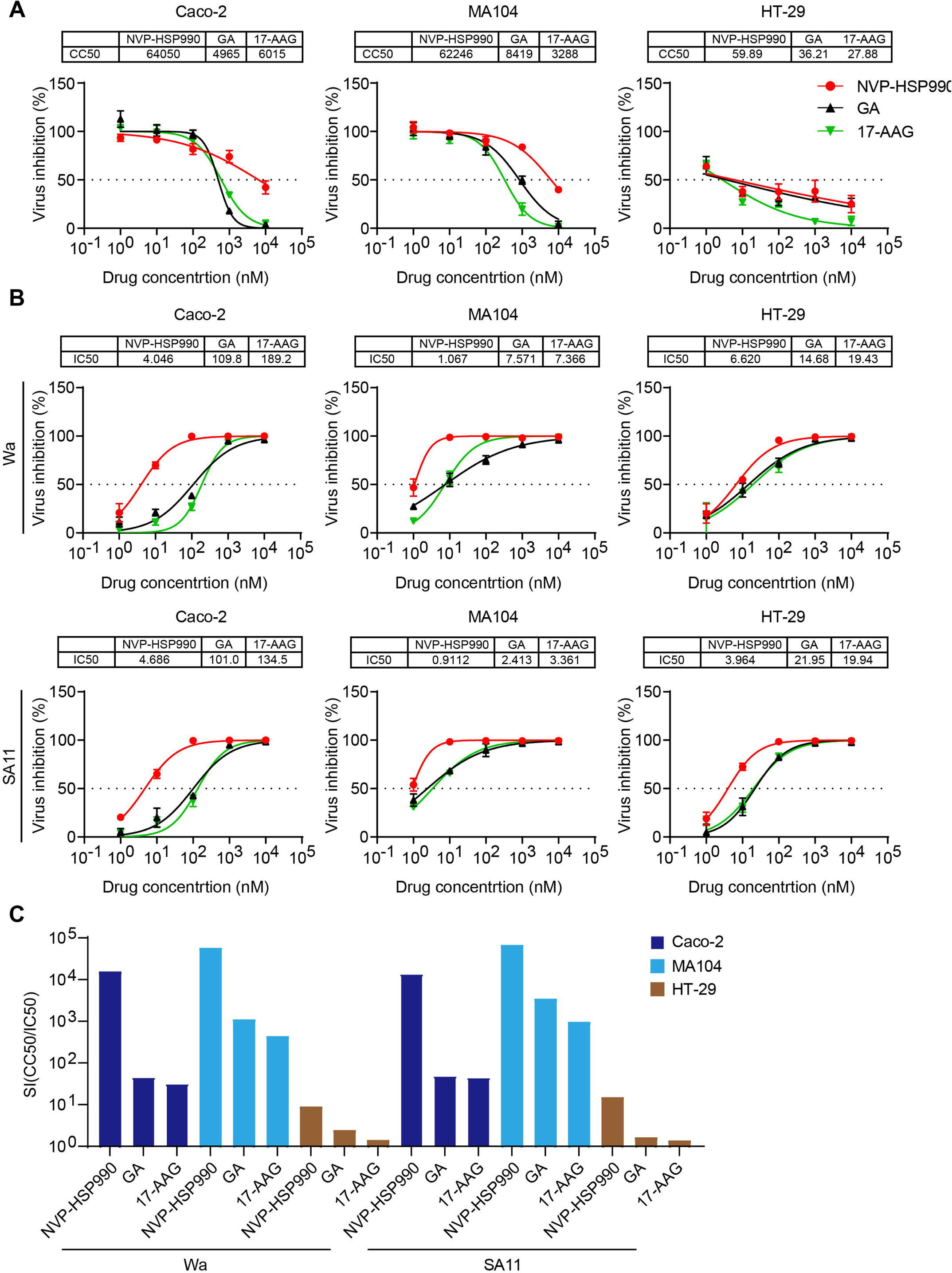
Cytotoxicity and inhibitory effects on RV reproduction of different HSP90 inhibitors. (A) 1X10^4^ Caco-2, MA104, or HT-29 cells were seeded in 96-well plates and cultured for 12 h, treated with different concentrations of NVP-HSP990, GA, or 17-AAG, and cell viability was evaluated by CCK-8 24 h later. CC50 values of drugs were calculated by the GraphPad Prism 9.0 software. (B) Caco-2, MA104, or HT-29 cells with 90% cell confluency in 96-well plates were infected with RV Wa or SA11 strains (MOI=1) and treated with indicated concentrations of NVP-HSP990, GA, or17-AAG. At 24 h p.i., viral reproduction was tested by PFA, and the IC50 value of each drug was calculated with the GraphPad Prism 9.0 software. (C) Selectivity index (SI) of NVP-HSP990, GA, and 17-AAG on different RV-infected cells. Data are presented as mean ± s.e.m. The experiments were performed in triplicate and the data are representative of two independent experiments.

To further validate the anti-RV effect of NVP-HSP990, Caco-2 cells were infected with RV Wa or SA11 strain with a series (0.001∼10) of MOI and cultured in DMEM containing 100 nM NVP-HSP990 for 24 h. We found that NVP-HSP990 significantly inhibited RV reproduction in all MOIs (Fig. 2A). We then evaluated the anti-RV effects of NVP-HSP990 which was administrated at different phases of RV infection. We found that administration of NVP-HSP990 before RV infection largely did not disturb the establishment of RV infection and RV reproduction (Supplementary Fig. 2A-C, Fig. 2B); NVP-HSP990 administration during RV infection only decreased RV reproduction to some distance (about 1 fold), indicating that NVP-HSP990 did not efficiently block RV cell entry (Fig. 2B). However, NVP-HSP990 administration after RV infection achieved excellent inhibitory effect on RV reproduction (over 100 folds); to be noted, the duration of action of NVP-HSP990 was important for its inhibitory effect, and the longer the duration of action of NVP-HSP990, the better the inhibitory effect on RV reproduction (Fig. 2B, C). qPCR analysis revealed significant reduction of genomic RNA synthesis of RV structural proteins VP2 and VP6 in RV-infected Caco-2 cells cultured with NVP-HSP990 (Fig. 2D), indicating that NVP-HSP990 significantly inhibited the transcription of RV antigens. Accordingly, expression of RV structural proteins VP6 and VP7 were remarkably reduced in RV-infected Caco-2 or MA104 cells treated with NVP-HSP990 (Fig. 2E). IFA also showed significant inhibition of viral antigens in RV-infected Caco-2 cells treated with 100 nM NVP-HSP990 (Fig. 2F).

**Fig. 2.**
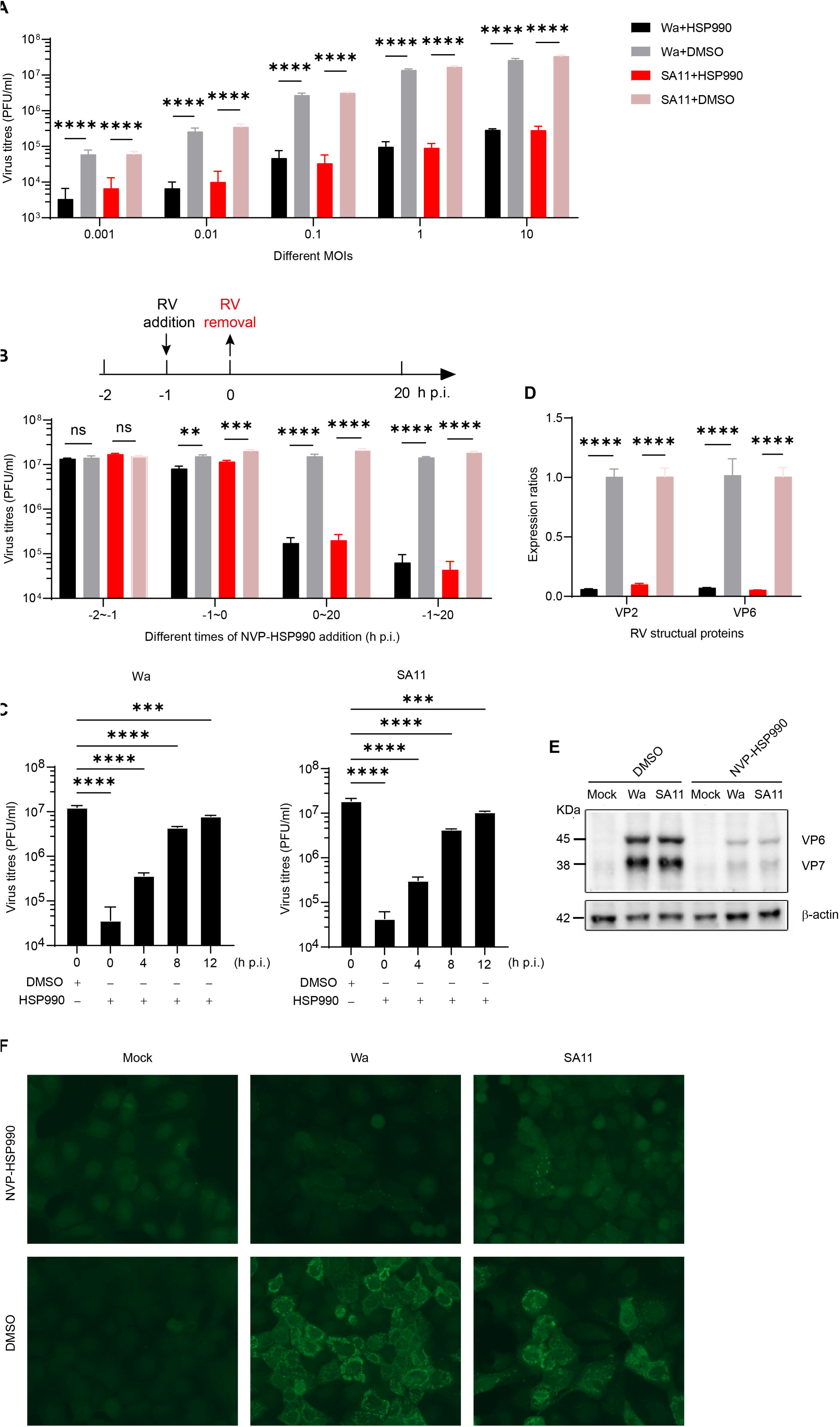
NVP-HSP990 significantly inhibited RV reproduction, viral gene transcription and viral antigen expression. (A) Caco-2 cells with 90% cell confluency in 96-well plates were infected with different MOIs (0.001∼10) of RV Wa or SA11 strains, and further cultivated with DMEM containing 100 nM NVP-HSP990 or equal amount of DMSO as control for 24 h, then viral production was tested with PFA. (B) Caco-2 cells with 90% cell confluency in 96-well plates were infected with RV Wa or SA11 strains (MOI=1). The cells were treated with 100 nM NVP-HSP990 before infection (-2∼-1 h p.i.), during infection (-1∼0 h p.i.), after infection (0∼20 h p.i.), or after virus exposure (-1∼20 h p.i.), or with the same volume of DMSO as control. Virus production was tested with PFA at 20 h p.i.. (C) Caco-2 cells with 90% cell confluency in 96-well plates were infected with RV Wa or SA11 strains (MOI=1). The cells were treated with 100 nM HSP990 at 0, 4, 8, 12 h p.i. or treated with the same volume of DMSO at 0 h p.i.. Virus production was tested with PFA at 24 h p.i.. (D) Caco-2 cells were infected with RV Wa or SA11 strains (MOI=1) and further cultivated with DMEM containing 100 nM NVP-HSP990 or equal volume of DMSO as control for 18 hours. Then the infected cells were harvested for qPCR analysis of RV structural protein VP2 and VP6. (E) Caco-2 cells were mock infected with PBS or infected with RV Wa or SA11 strains (MOI=1) and further cultivated with DMEM containing 100 nM NVP-HSP990 or equal volume of DMSO as control for 18 hours. Then the infected cells were harvested for WB analysis expression of RV structural protein VP6 and VP7. (F) Caco-2 cells growing on coverslips were mock infected with PBS or infected with RV Wa or SA11 strains (MOI=1), and cultivated with DMEM containing 100 nM NVP-HSP990 or equal volume of DMSO as control after infection for another 18 hours. Then the infected cells were applied for immune staining of RV antigens (green) and images were captured using Zeiss Axio Imager A2. Data are presented as mean ± s.e.m (A-D). The experiments were performed in triplicate (A-D) and the data are representative of two independent experiments (A-F). ns: no significance, \*\**P* < 0.01, \*\*\**P* < 0.001, \*\*\*\**P* < 0.0001 (two-way ANOVA test (A, B, D) and one-way ANOVA test (C)).

### 3.2 NVP-HSP990 alters the life state of host cells

To find out the effects of NVP-HSP990 on the impacts to host cells caused by RV infection, Caco-2 cells were mock infected with PBS or infected with RV Wa or SA11 strains (MOI=3) and further cultivated with DMEM containing 100 nM NVP-HSP990 or equal volume of DMSO as control for 24 h. Then, the infected cells were harvested for RNAseq analysis. PCA analysis indicated that RV infection and administration of NVP-HSP990 were important factors influencing gene expression patterns of host cells (Fig. 3A). As expected, RV infection (both Wa and SA11 strains) caused significant transcriptional changes in host cells, while NVP-HSP990 remarkably reduced the influence of RV infection on gene transcription in host cells (Fig. 3B). To be noted, although most KEGG pathway alterations caused by RV infection were eliminated by NVP-HSP990 administration, some pathways such as IL-17, TNF singling pathways, which are related to innate immunity or inflammation, remained altered even in the presence of NVP-HSP990 (Fig. 3C-F). Next, we analyzed the direct impacts of NVP-HSP990 host cells. We found that NVP-HSP990 caused significant transcriptional alterations of various genes in both mock- and RV-infected host cells (Fig. 4A), and these alterations were mainly related to cell cycle, DNA replication, and various cancer-, signaling- or metabolism-related pathways in host cells (Fig. 4B). Among these genes, 112 genes were up-regulated and 287 genes were down-regulated by NVP-HSP990 in all mock-, Wa-, and SA11-infected host cells (Fig. 4C, Supplementary Table 2, 3). These up-regulated genes were mainly related to KEGG pathways like protein processing in endoplasmic reticulum, antigen processing and presentation, etc., and the down-regulated genes were mainly related to cell cycle, DNA replication, and various cancer related KEGG pathways (Supplementary Fig. 3A, C). To be noted, 75 genes were up-regulated and 96 genes were down-regulated by NVP-HSP990 in both Wa-, and SA11-infected but not in mock-infected host cells (Fig. 4C, Supplementary Table 4, Supplementary Table 5), which were mainly related to metabolism- or inflammation-related KEGG pathways respectively (Supplementary Fig. 3B, D).

**Fig. 3.**
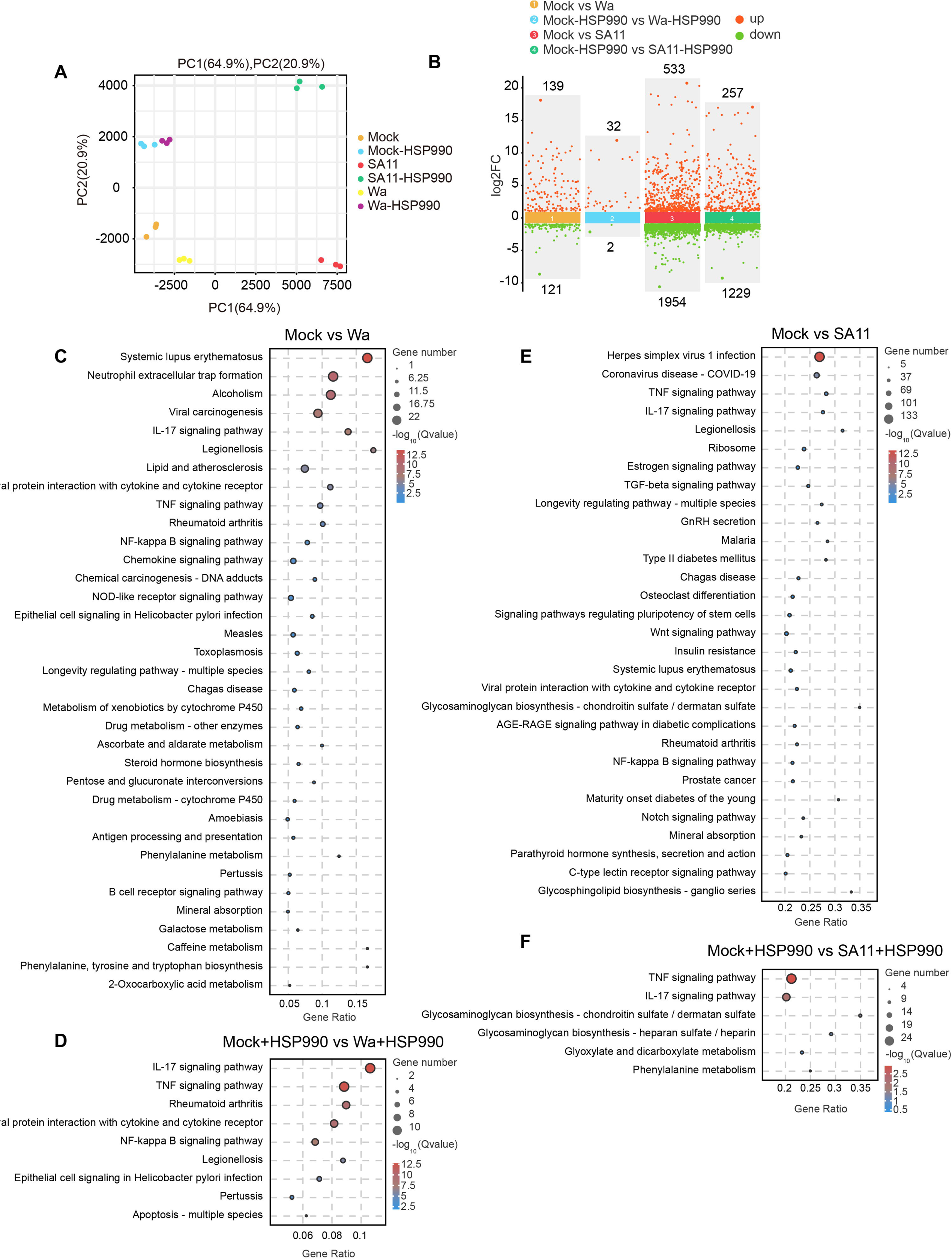
NVP-HSP990 eliminates the effects of RV infection on host cell signaling pathways. Caco-2 cells were mock infected with PBS or infected with RV Wa or SA11 strains (MOI=3) and further cultivated with DMEM containing 100 nM HSP990 or equal volume of DMSO as control for 24 hours. Then, the infected cells were harvested for RNAseq analysis. PCA analysis (A), multiple differential scatter plots of compared groups (B), and bubble charts of differential KEGG pathways in compared groups (C-F) are shown.

**Fig. 4.**
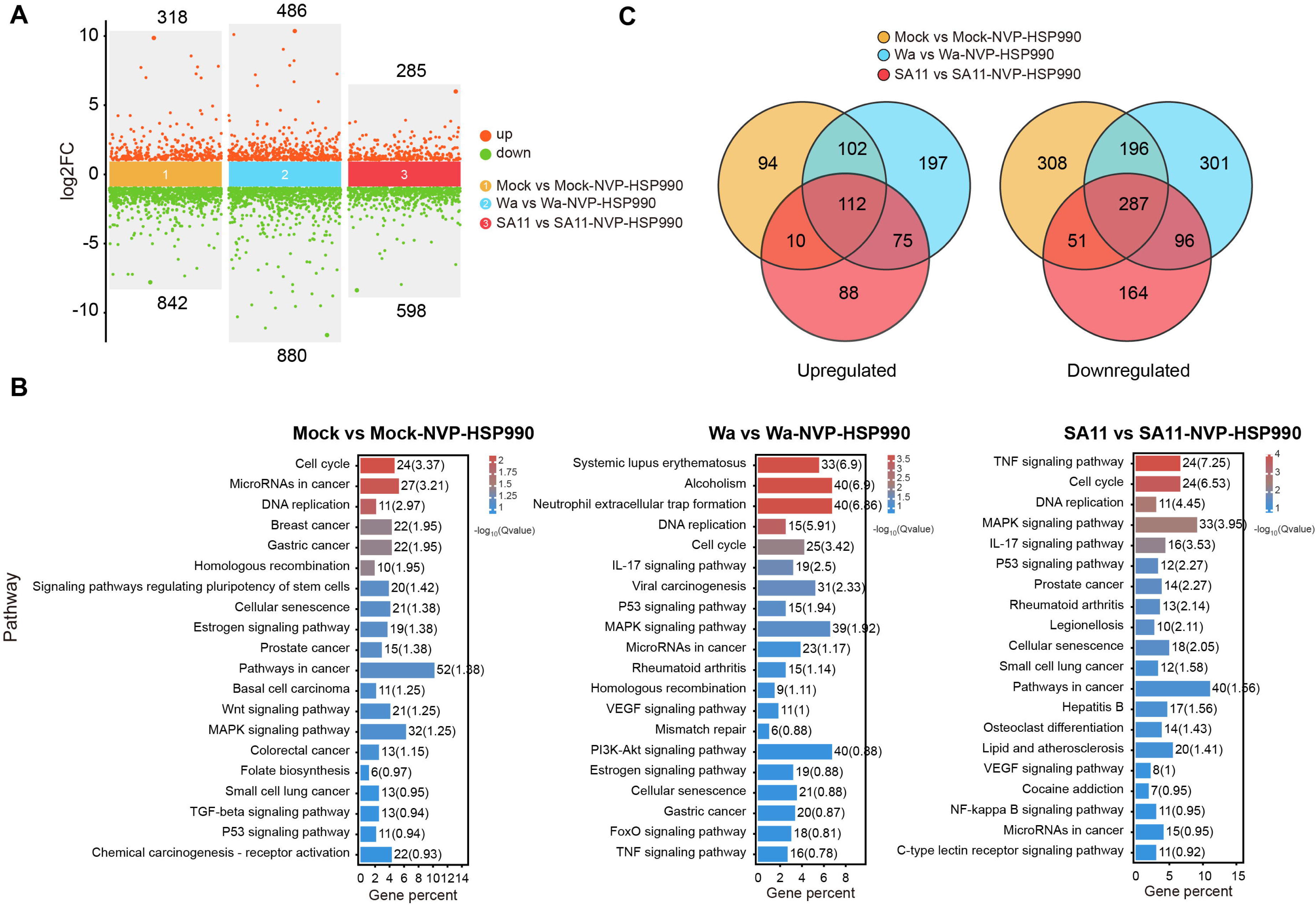
NVP-HSP990 alters the life state of host cells. Caco-2 cells were mock infected with PBS or infected with RV Wa or SA11 strains (MOI=3) and further cultivated with DMEM containing 100 nM NVP-HSP990 or equal volume of DMSO as control for 24 h. Then, the infected cells were harvested for RNAseq analysis. Multiple differential scatter plots of compared groups (A), bar charts of differential KEGG pathways between compared groups (B), and Venn diagrams of up- and down-regulated genes between compared groups (C) are shown.

### 3.3 NVP-HSP990 modulated RV-induced activation of MAPK signaling pathway in Caco-2 cells

MAPK signaling pathway is mainly composed of ERK, JNK, and p38 signals and usually plays important roles in viral infections and host antiviral immunity(29, 30). We found that NVP-HSP990 significantly up-regulated some but down-regulated more MAPK-related genes (Fig. 5A, Supplementary Fig. 3A, C). The up-regulated MAPK-related genes included some heat shock proteins (HSPB1, HSPA1A, HSPA1B, etc.) and some cytokine/cytokine receptors (PGF, PDGFRA, PDGFRB, etc.), while the down-regulated genes contained JUN, JUND, MYC, DUSP2, DUSP5, etc., which are involved in signal transduction (Fig. 5A, Supplementary Fig. 4, Supplementary Table 6-8), indicating that NVP-HSP990 mainly inhibited MAPK signaling pathway in RV infection. In fact, MAPK signaling pathway is activated and critical for RV reproduction(18, 31, 32). We found that NVP-HSP990 robustly inhibited the activation of ERK, JNK, and p38 signals in RV-infected Caco-2 other than MA104 cells, though RV infection generated significant activation of these MAPK signaling pathways in both cell lines (Fig. 5B). These results indicated that NVP-HSP990 might specifically inhibited MAPK activation in intestinal epithelial cells which are natural target cells of RV, thus favoring its anti-RV effect *in vivo*. Tight junctions are crucial for the survival and function of mature intestinal epithelial cells, and destruction of tight junctions causes intestinal inflammation and disorder(33, 34). The formation of tight junctions is modulated by intracellular signaling pathways including MAPK signaling pathway(35, 36). As we found NVP-HSP990 significantly inhibited MAPK signaling pathway in Caco-2 cells, accordingly, it was also found that NVP-HSP990 effectively alleviated RV-induced disruption of tight junctions of Caco-2 cells (Fig. 5C).

**Fig. 5.**
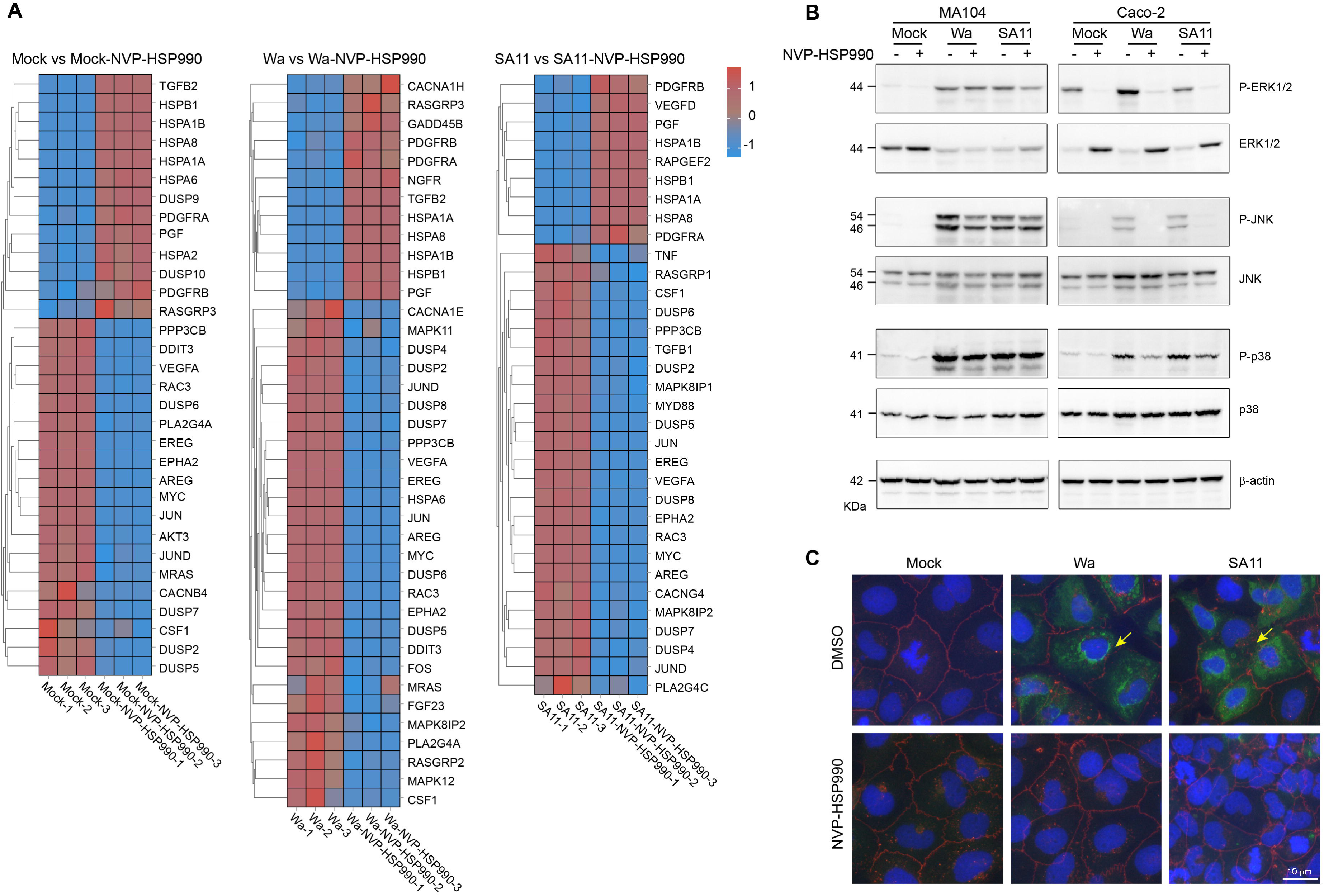
NVP-HSP990 inhibited RV-induced activation of MAPK signaling pathway and prevented disruption of tight junction in Caco-2 cells. (A) Heat maps of differentially expressed genes belonging to MAPK KEGG pathway. (B) MA104 and Caco-2 cells were infected with RV Wa or SA11 strains (MOI=1) or mock infected with PBS, and then treated with 100 nM HSP990 (+) or equal volume of DMSO as control (-) for 20 h. Then the infected cells were harvested for WB analysis of expression of MAPK components (JNK, Phospho-JNK (P-JNK), p38, Phospho-p38 (P-p38), ERK1/2, and Phospho-ERK1/2 (P-ERK1/2)) with β-actin as internal reference. (C) Caco-2 cells growing on coverslips were mock infected or infected with RV Wa or SA11 strains (MOI=1), and cultivated with DMEM containing 100 nM NVP-HSP990 or equal volume of DMSO as control after infection for another 18 h. Then the infected cells were applied for immune staining of RV antigens (green) and ZO-1 (red) which is marker of tight junction, as well as DAPI staining of nucleus (blue). Disruption of tight junction is shown by yellow arrows. Data are representative of two independent experiments.

### 3.4 NVP-HSP990 alleviated RV infection in suckling mice

As NVP-HSP990 possessed potent anti-RV effects *in vitro* as proved above, we wondered whether NVP-HSP990 was effective on control of RV diarrhea. To this end, 7-days-old BALB/c suckling mice (with a body weight of 2.5∼4.8 g) were infected with 1 × 10^6^ PFU RV SA11 strain orally. Different doses of NVP-HSP990 were orally administrated at 2 h p.i., and diarrhea scores of the infected suckling mice were evaluated everyday as described(18, 28). At 24 h p.i. (day 1), it was shown that average diarrhea scores reduced along increasing doses of NVP-HSP990, and the reduction was especially remarkable at doses of 100∼1000 μg/Kg (Fig. 6A). NVP-HSP990 inhibited RV diarrhea occurrence with an EC50 value of 142.3 μg/Kg, and reduced diarrhea score with an EC50 value of 135.5 μg/Kg (Fig. 6B, C). Through continuous monitoring, we found that treatment with 1 mg/Kg NVP-HSP990 did not hinder body growth of RV-infected suckling mice but significantly alleviated RV diarrhea (Fig. 6D-F). Therefore, these results indicated that NVP-HSP990 had a potent ability to alleviate RV diarrhea.

**Fig. 6.**
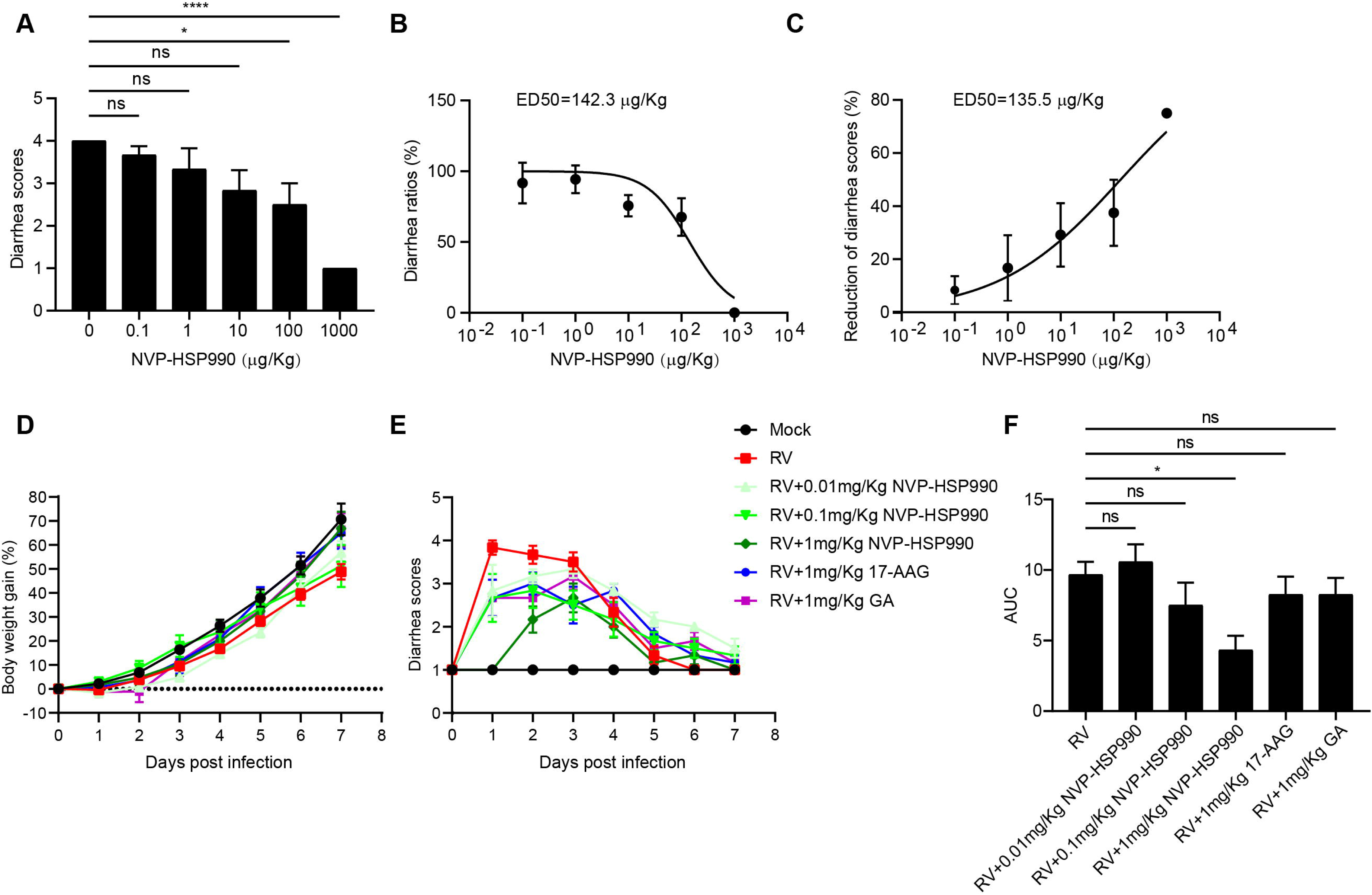
NVP-HSP990 inhibited RV diarrhea in suckling mice. (A-C) 7-days-old BALB/c suckling mice were orally inoculated with 1×10^6^ PFU RV SA11 strain. At 2 h p.i., the mice were orally treated with a series dose (0, 0.1, 1, 10, 100, 1000 μg/Kg) of NVP-HSP990 (n=6 mice /group) and diarrhea scores of the RV-infected suckling mice at 24 h p.i. are recorded and analysed (A); the ED50 values of NVP-HSP990 to diarrhea occurrence (B) or diarrhea score (C) at 24 h p.i. were calculated by the GraphPad Prism 9.0 software. (D-F) 7-days-old BALB/c suckling mice were orally inoculated with 1×10^6^ PFU RV SA11 strain or mock infected with PBS. At 2 h p.i., the RV-infected suckling mice were treated with a series doses (0.01, 0.1, and 1 mg/Kg) of NVP-HSP990, 1 mg/Kg 17-AAG, or 1 mg/Kg GA respectively, with equal volume of DMSO as control (n=6 mice/group). Body weight gains (D) and diarrhea scores (E) of the suckling mice were monitored from 0 to 7 days post infection, and area under curve of diarrhea scores in each group was assessed by GraphPad Prism 9.0 software with the curve of mock-infection was set as baseline (F). Data are presented as mean ± s.e.m and are representative of two independent experiments. ns: no significance, \**P* < 0.05, \*\*\*\**P* < 0.0001 (one-way ANOVA test (A, F)).

As NVP-HSP990 was competent in control of RV diarrhea as shown above, we wondered whether it also significantly inhibited RV replication *in vivo* as it did *in vitro*. To this end, RV-infected BALB/c suckling mice were treated with 1 mg/Kg NVP-HSP990 at 2 h p.i. with equal amount of DMSO as control. At 16 h p.i., infectious virus contents and viral antigens in intestines (including Duodenum, Jejunum, and Ileum) were analysed. We found that infectious RV contents and viral antigens in Jejunum and Ileum were remarkably reduced in RV-infected mice when treated with NVP-HSP990, though there was no significant difference in duodenum (Fig. 7A, B). Accordingly, transcription of RV structural protein VP6 was remarkably reduced in epithelial cells of Jejunum and Ileum from NVP-HSP990-treated RV-infected mice (Fig. 7C). As RV infection usually leads to obvious pathological changes to Ileum, we checked the protective effect of NVP-HSP990 on Ileum lesions caused by RV infection. NVP-HSP990 showed no obvious pathological effect on Ileum tissue in mock-infected mice, but it obviously prevented pathological lesions caused by RV infection (Fig. 7D). Therefore, these results indicated that NVP-HSP990 competently inhibited RV replication and prevented pathological lesions in intestine caused by RV infection.

**Fig. 7.**
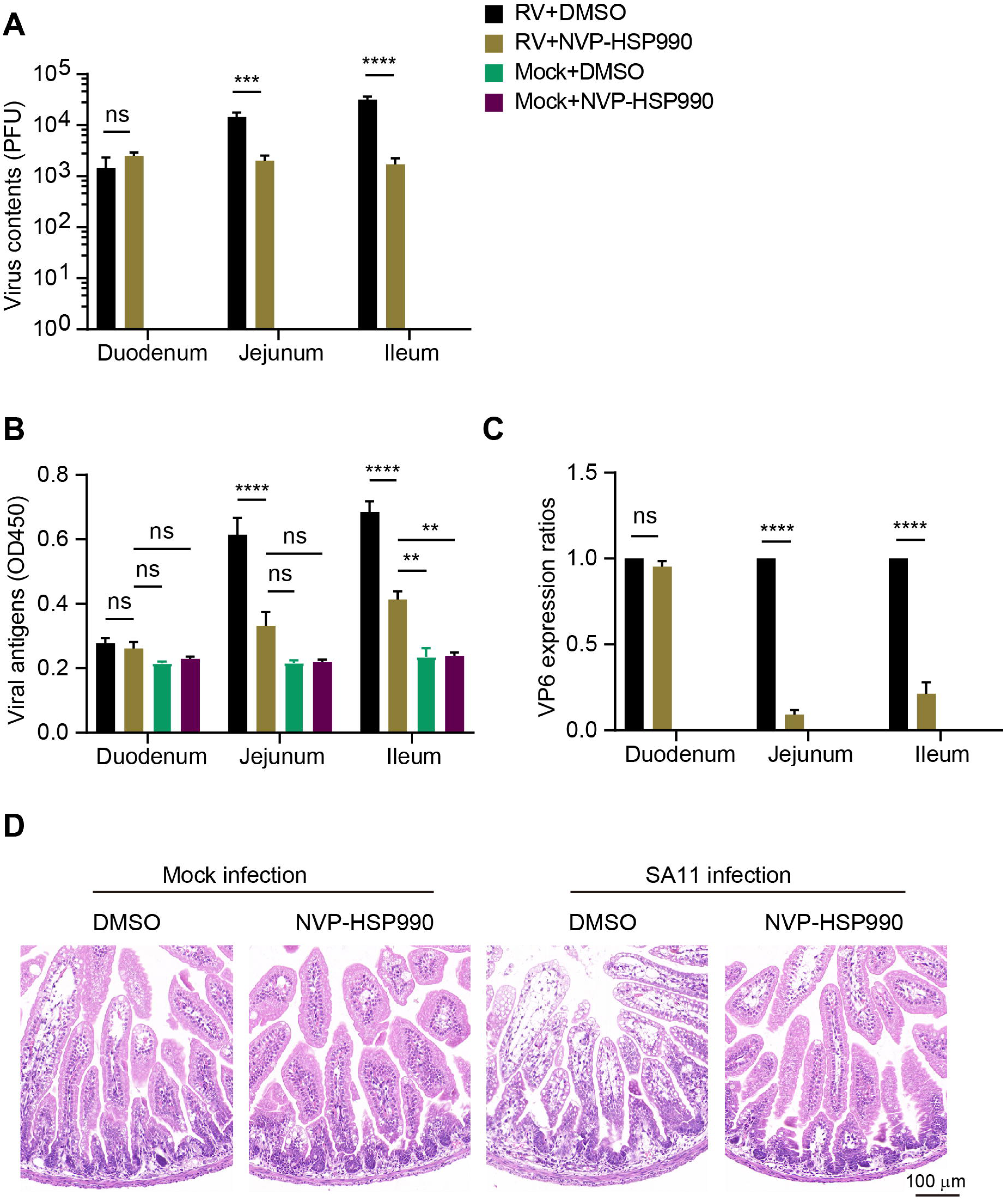
NVP-HSP990 inhibited RV infection and alleviated pathological changes in the intestine of suckling mice. (A-C) 7-days-old BALB/c suckling mice were orally inoculated with 1×10^6^ PFU RV SA11 strain or mock-infected with PBS. At 2 h p.i., the mice were orally treated with 1 mg/Kg of HSP990 with equal volume of DMSO as control (n=3 mice/group). Then, the infectious RV particles and RV antigens in contents of intestines (Duodenum, Jejunum, and Ileum) were analyzed with PFA (A) and ELISA (B) respectively at 16 h p.i., and transcription of RV structural protein VP6 in intestinal epithelial cells was analysed with qPCR (C). (D) Ileum of the suckling mice were fixed in 4% paraformaldehyde and subjected to paraffin section, followed by staining with hematoxylin and eosin. Data are presented as mean ± s.e.m (A-C) and are representative of two independent experiments (A-D). ns: no significance, \*\**P* < 0.001, \*\*\**P* < 0.001, \*\*\*\**P* < 0.0001 two-way ANOVA test).

## 4. Discussion

Cell stress caused by viral infection results in enhanced expression of various heat shock proteins, including HSP40, HSP70, and HSP90 etc., and viral replications have been shown to be directly or indirectly dependent on one or more of these HSPs(37, 38). Various reports revealed that many viruses, including RV, require the participation of HSP90 in their replication, for gene knockdown or the use of chemical inhibitors of HSP90 could significantly inhibit viral replications *in vitro*(10–12, 17, 39, 40). Therefore, it sounds promising to develop novel HSP90-targeting, broad-spectrum antiviral drugs, which would be more stable in antiviral effect and less affected by viral mutation. Unfortunately, considering their antiviral efficacy and toxicity, there has been no HSP90 inhibitors in clinical use for viral control till now. In this study, we showed that NVP-HSP990, an orally available small-molecule inhibitor of HSP90, inhibited RV infection with high efficacy and low toxicity and significantly alleviated RV diarrhea in suckling mice, thus suggesting that targeting HSP90 remains to be a promising strategy for antiviral drug development.

Small-molecule inhibitors of HSP90 are currently popular and pursued as a potential candidates for treating tumors and viral infections(19, 41–43). The initial designs of small-molecule HSP90 inhibitors were mainly based on the mechanism of the natural HSP90 inhibitor GA or Radicicol, which targets the ATP pocket of N-terminal domain of HSP90 to block HSP90 ATPase activity, and the representatives include GA derivatives such as 17-allylamino-17-demethoxygeldanamycin (17-AAG), 17-N, N-dimethylaminoethyl-17-methoxygeldanamycin (17-DMAG), 17-AAG hydroquinone hydrochloride (IPI-504) and Radicicol derivatives such as NVPAUY922, AT13387, and Ganetespib. Recently, with the increasing understanding of HSP90 subtypes and the interactions between HSP90 and its client proteins, a series of novel HSP90 inhibitors have emerged, which selectively inhibit HSP90 subtypes or specifically block the interaction between HSP90 and client proteins, so as to improve their inhibitory efficacy and reduce their toxicity(44, 45). NVP-HSP990 is a novel small-molecule HSP90 inhibitor with much smaller molecular weight than GA and 17-AAG. NVP-HSP990 targets the N terminals of both HSP90α and HSP90β with high oral bioavailability and was previously tested in treatment of tumors and Huntington’s disease(24, 46–48). However, there have been few experimental reports on the role of NVP-HSP990 in controlling viral infections, though a recent in silico study predicted that NVP-HSP990 might alleviate COVID-19 symptoms through anti-inflammatory effects on SARs-CoV-2 infected lung cells(49). Here, our data showed that NVP-HSP990 did not hinder the establishment of initial RV infection, meaning that NVP-HSP990 did not interfere with RV membrane attachment, entry or disruption of the outer capsid to yield DLP (the active transcription unit of RV) for transcription initiation; however, NVP-HSP990 potently inhibited RV replication with much bigger SI than traditional HSP90 inhibitors such as GA and 17-AAG. Therefore, our findings implied that NVP-HSP990 may be a promising antiviral candidate for RV infection.

To remodel host cell’s life system for better viral replication, viral infection usually triggers the modulation of host signaling pathways, resulting in transcriptional changes of various host genes(50–52). On the other hand, viral infection inevitably triggers the activation of inflammation-associated signaling pathways for host antiviral innate immunity. In this study, RV infection triggered significant transcriptional changes of genes involved in various pathways of cell life; however, in the presence of NVP-HSP990, the impact of RV infection on most pathways faded away. To be noted, the impact of RV infection on inflammation-associated signaling pathways (e.g., IL-17, TNF signaling pathways) were still significant even in the presence of NVP-HSP990, suggesting that NVP-HSP990 could not significantly block the activation of host antiviral innate immunity during RV infection though it potently inhibited RV replication, which would be a great advantage of NVP-HSP990 to be an antiviral candidate.

Upon NVP-HSP990 treatment, the most affected pathways in Caco-2 cells included cell cycle, DNA repair and cancer-related pathways, which is consistent with its role as an effective anti-tumor drug. To be noted, the MAPK signaling pathway was negatively regulated by NVP-HSP990 in both non-infected and RV-infected Caco-2 cells, suggesting that the MAPK signaling pathway is sensitive to HSP90 inhibition by NVP-HSP990, which is in accordance with the report that proper folding of RAF kinases of the RAS-MAPK signaling pathway relies on the interaction with HSP90(53). RV infection induces the activation of most MAPK signaling pathways, including ERK, JNK and p38, among which p38 and ERK are critical for RV replication, while JNK is critical for host IFN-β production(31, 32, 54, 55). Here we showed that NVP-HSP990 robustly inhibited RV-induced activation of ERK, JNK and p38 in Caco-2 cells other than MA1014 cells, suggesting that NVP-HSP990 could inhibit RV replication by specifically blocking MAPK signaling activation in intestinal cells, which should facilitate its anti-RV efficacy *in vivo*. Disruption of tight junctions between intestinal epithelial cells leads to the necrosis or anoikis of intestinal cells, which is usually a key step in the pathogenesis of intestinal infectious diseases(56). Many studies show that activation of the p38MAPK signaling pathway disrupts cellular tight junctions(36, 57). In this study, RV infection significantly activated p38MAPK signaling and disrupted cellular tight junctions in Caco-2 cells, which were consistent with previous studies(32, 58). Nevertheless, NVP-HSP990 significantly inhibited the activation of MAPK signaling pathway, thus resulted in significant alleviation of the disruption of tight junction in RV-infected Caco-2 cells.

Besides gastrointestinal symptoms such as diarrhea and vomiting, severe RV infections often impinge important organs such as the brain, heart, kidneys, and liver. For example, RV infection usually causes epileptic seizures in children, although the mechanism is currently unclear(5). Conventional antiviral drugs such as nucleoside analogues and type I interferon are usually not included in the treatment of RV diarrhea due to adverse side effects(6, 7), thus symptomatic treatment is usually the main choice, which sometimes leads to systemic RV infection and even mortality due to persistent RV infection. Therefore, the treatment of RV infection should not merely focus on symptomatic treatment, timely antiviral treatment is also important to reduce complications and child mortality. Here, we demonstrated that NVP-HSP990 effectively alleviated RV diarrhea of BALB/c suckling mice and competently inhibited RV replication and prevented pathological lesions in intestine caused by RV infection. To be noted, NVP-HSP990 is able to penetrate the blood-brain barrier and extremely low dose of NVP-HSP990 has significant therapeutic effects on epileptic seizures (25). Therefore, NVP-HSP990 might be a promising candidate of antiviral drugs especially suitable for RV infection which usually offense the brain.

## 5. Conclusion

In this study we demonstrated that a small-molecule HSP990 inhibitor NVP-HSP990 was a novel potent antiviral agent with much higher SI compared to traditional HSP990 inhibitors. NVP-HSP990 robustly blocked RV replication with low cytotoxicity *in vitro*, repressed RV-induced activation of MAPK signaling pathway and disruption of tight junctions in intestinal cells, effectively alleviated RV diarrhea of suckling mice, competently inhibited RV replication, and obviously prevented pathological lesions in intestine caused by RV infection. These findings indicated that NVP-HSP990 can be a novel promising candidate of antiviral drugs for alleviating RV infection.

## Abbreviations used in this paper

17-AAG: 17-allylamino-17-demethoxygeldanamycin
17-DMAG: N-dimethylaminoethyl-17-methoxygeldanamycin
ANOVA: analysis of variance
ATCC: American type culture collection
CCK-8: cell counting kit-8
CC50: 50% cytotoxic concentration
COVID-19: Corona Virus Disease 2019
DAPI: 2-(4-Amidinophenyl)-6-indolecarbamidine dihydrochloride
DMEM: Dulbecco’s modified eagle medium
DMSO: dimethyl sulfoxide
DTT: dithiothreitol
EC50: concentration for 50% of maximal effect
ELISA: enzyme-linked immunosorbent assay
ERK: extracellular signal-regulated kinase
FBS: fetal bovine serum
FITC: fluorescein 5-isothiocyanate
FPKM: fragment per kilobase of transcript per million mapped reads
GA: geldanamycin
GRP94: glucose-regulated protein 94
HSP90: heat shock protein 90
IC50: half maximal inhibitory concentration
IFA: immunofluorescence assay
IL-17: interleukin 17
JNK: c-Jun N-terminal kinase
KEGG: Kyoto Encyclopedia of Genes and Genomes
MAPK: mitogen-activated protein kinase
MOI: multiplicity of infection
p38: p38 mitogen-activated protein kinase
PFA: plaque formation assay
PFU: plaque formation unit
qPCR: quantitative Real-time PCR
RV: rotavirus
RNA-seq: RNA-sequencing
SI: selectivity index
TNF: tumor necrosis factor
TRAP1: TNF receptor associated protein 1
WHO: world health organization.

## Acknowledgements

We thank Dr. Elschner (Friedrich-Loeffler-Institute) for gift of MA104 cells; Guangzhou Genedenovo Biotechnology for RNA-seq analysis; Servicebio for assistant of histopathology assay.

---

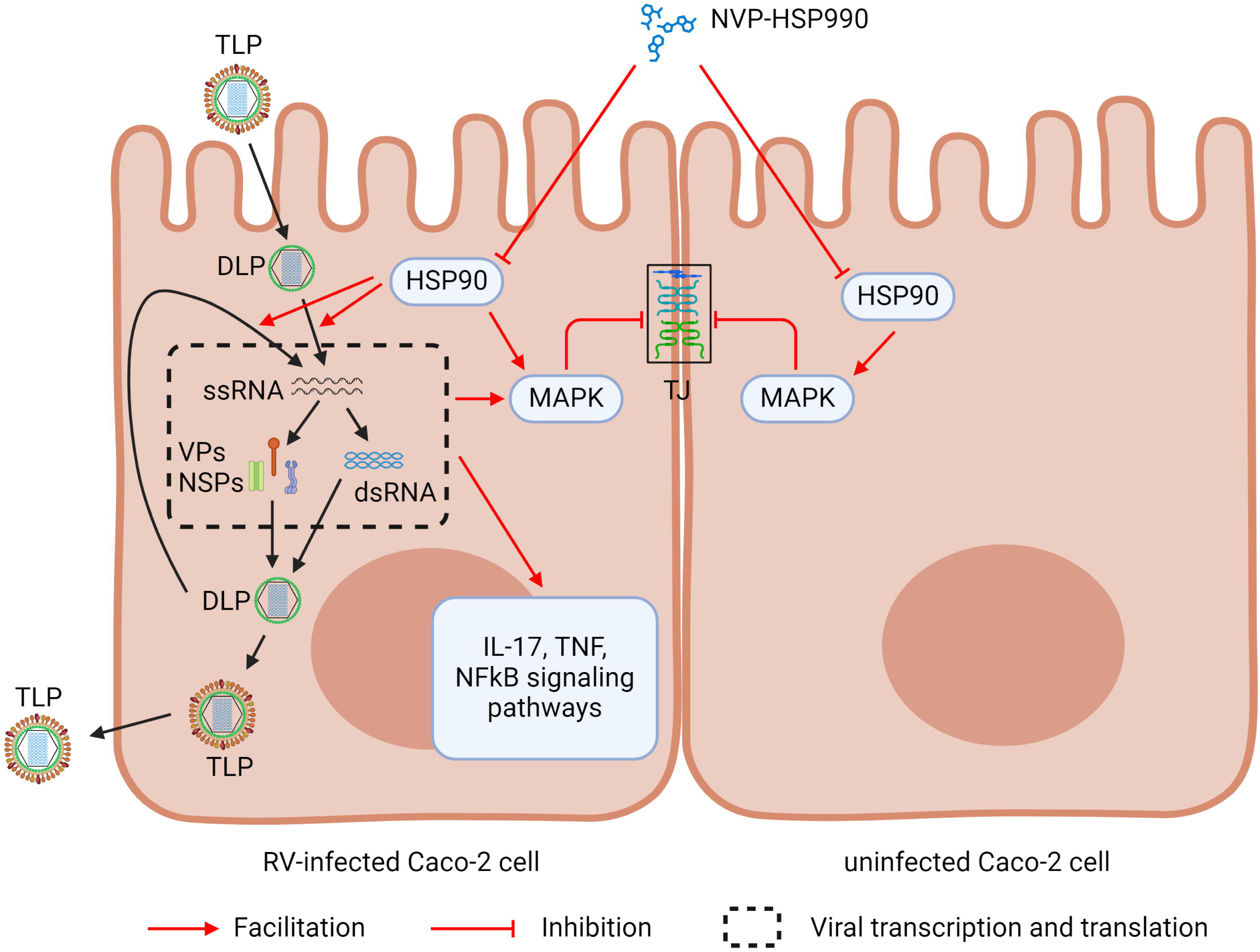

