## Supplementary Figure 1-4 for "HSP90 inhibitor NVP-HSP990 alleviates rotavirus infection"

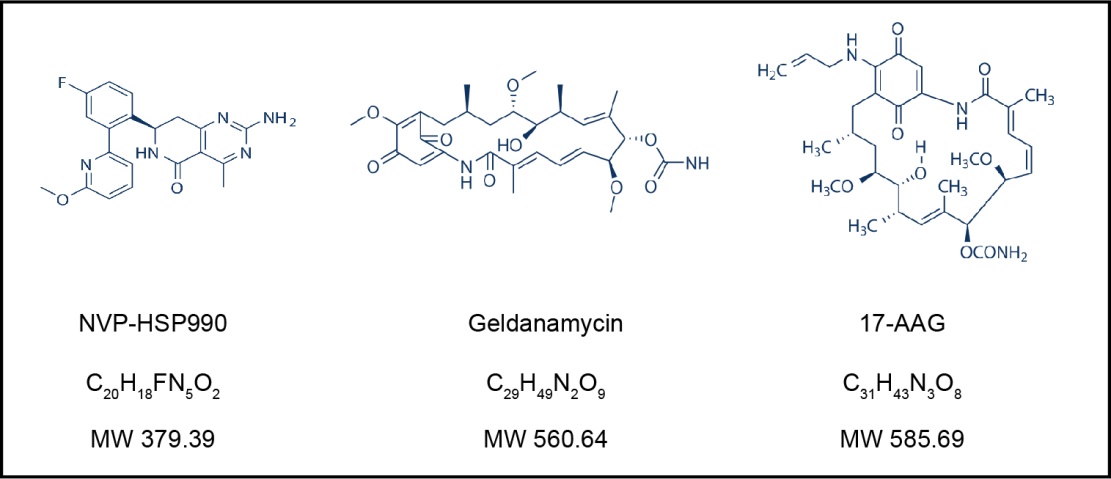

**Supplementary Fig. 1. Chemical structures of NVP-HSP990, Geldanamycin, and 17-AAG.**

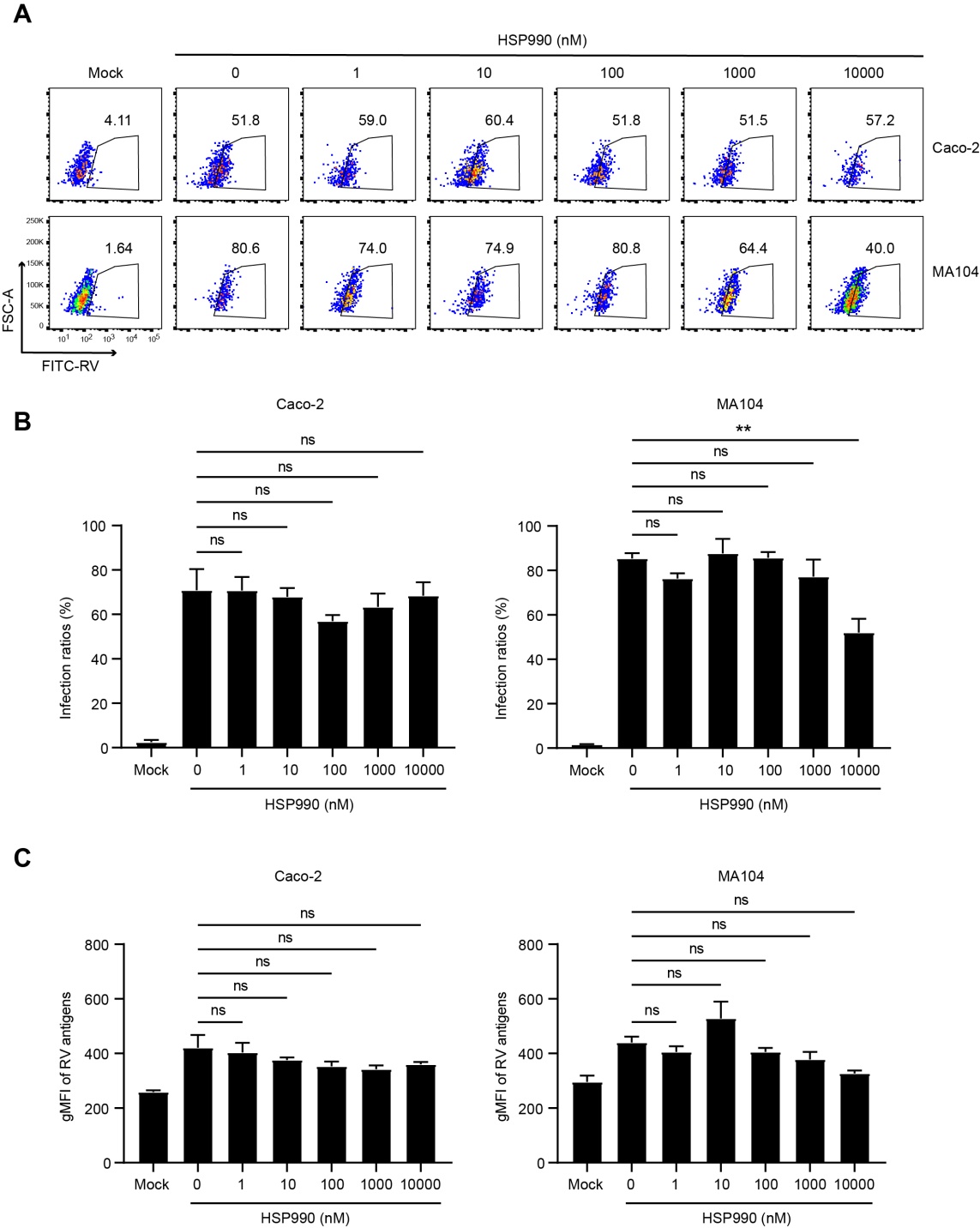

**Supplementary Fig. 2. HSP990 did not influence the establishment of RV infections.** Caco-2 or MA104 cells were treated with a series of NVP-HSP990 concentrations (0, 1, 10, 100, 1000, 10000 nM) for 2 h before RV Wa strain infection (MOI=1) respectively. After washing with 0.01 M PBS for 3 times, the cells were cultured in DMEM for 20 h and then harvested for FACS analysis of infection ratios and viral antigen expression. Representative count plots (A) and statistics of infection ratios (B) and expression of RV antigens (C) are shown. Data are presented as mean ± s.e.m and are representative of two independent experiments. ns: no significance, ***P* < 0.01

(one-way ANOVA test).

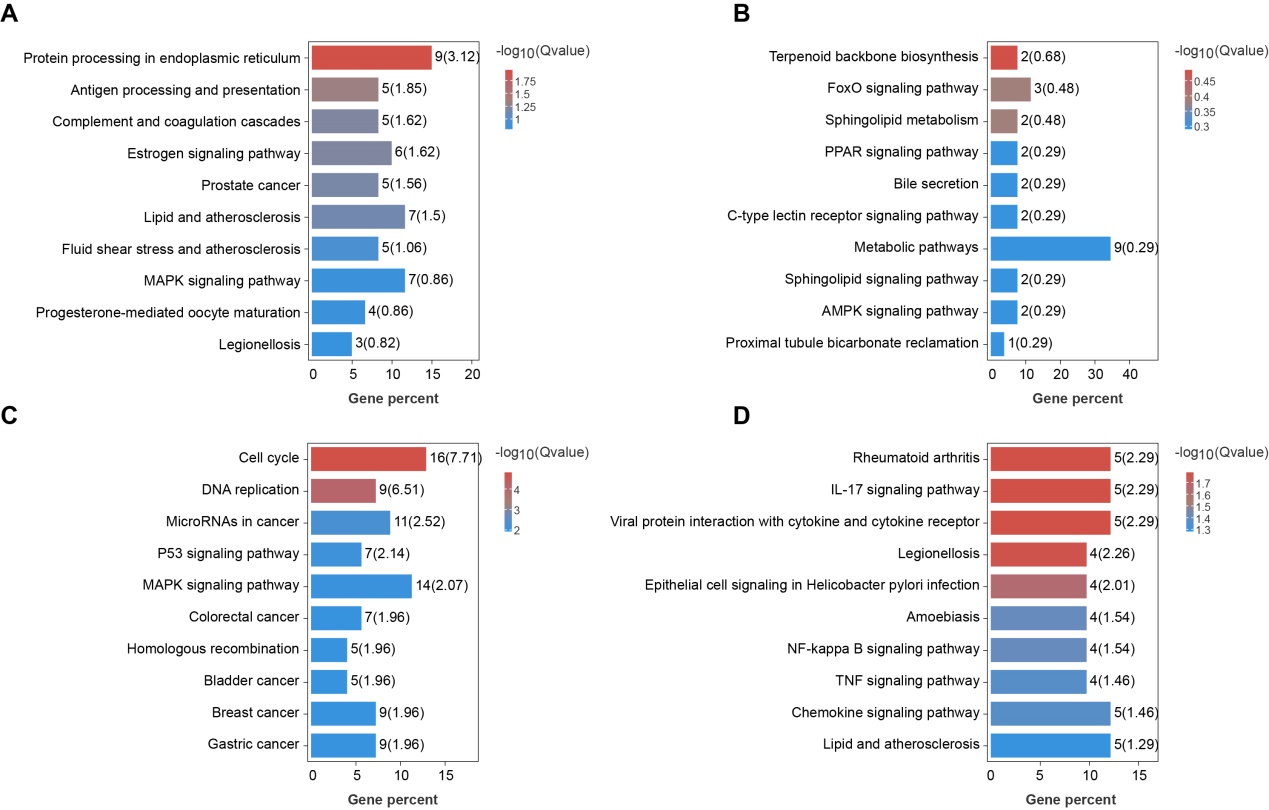

**Supplementary Fig. 3. NVP-HSP990 alters the life pathways of host cells.** Caco-2 cells were mock infected with PBS or infected with RV Wa or SA11 strains (MOI=3) and further cultivated with DMEM containing 100 nM NVP-HSP990 or equal volume of DMSO as control for 24 h. Then, the infected cells were harvested for RNAseq analysis. (A, C) Up-regulated (A) and down-regulated (C) KEGG pathways in all mock-, Wa- and SA11- infected Caco-2 cells. (B, D) Up-regulated (B) and down-regulated (D) KEGG pathways in both Wa- and SA11- infected but not in mock-infected Caco-2 cells.

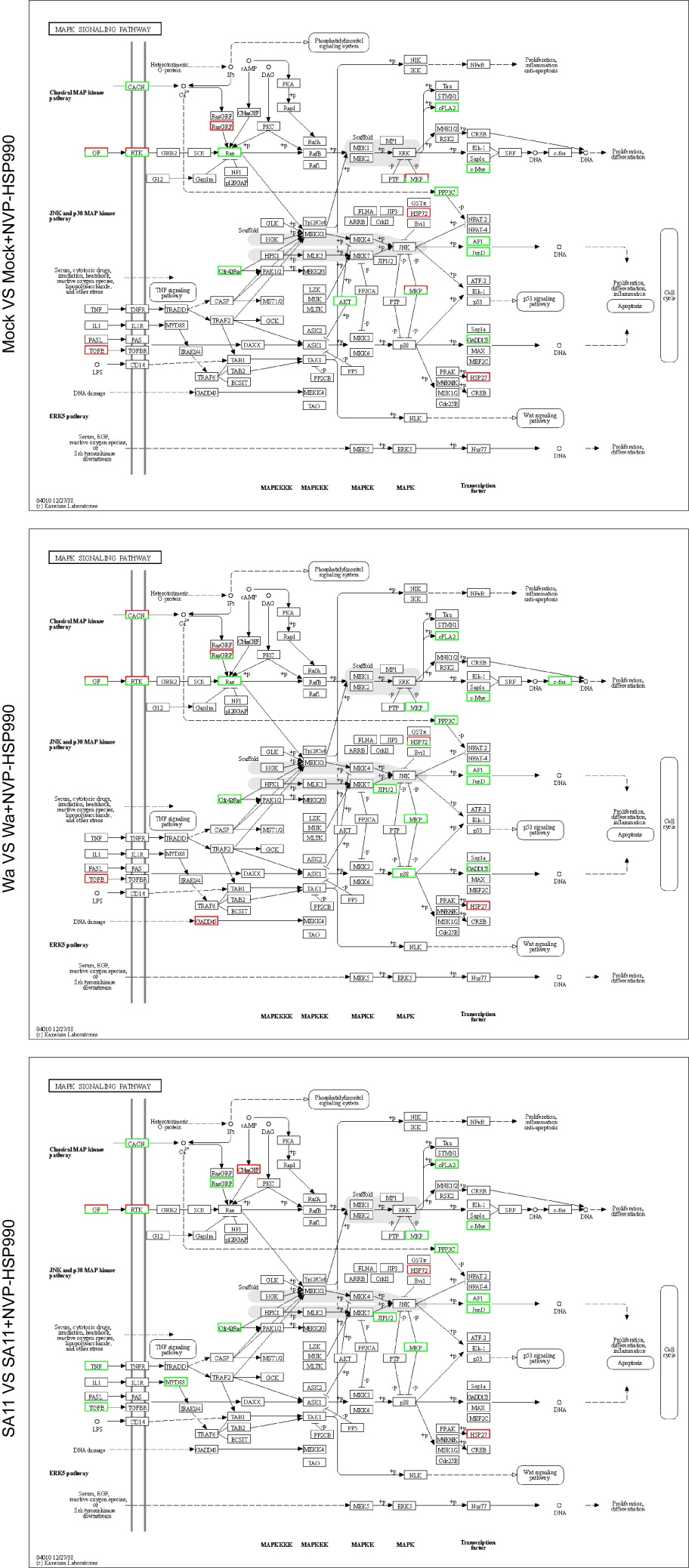

**Supplementary Fig. 4. Altered MAPK signaling pathway (KEGG) in mock- or RV- infected Caco-2 cells by NVP-HSP990. Red frames indicate up-regulated gene, while green frames indicate down-regulated gene.**
