## Supplementary Table 1 for "HSP90 inhibitor NVP-HSP990 alleviates rotavirus infection"

Supplementary Table 1. Primers used in qPCR analysis.

| Primer name | Forward primer (5’ – 3’) | Reverse primer (5’ – 3’) |
| --- | --- | --- |
| VP2 | ttgcttgcgttcgtttcagc | tggggttggcgtttacagtt |
| VP6 | tgctattaacgcaccagcca | aacctttccgcgtctggtag |
| Mouse β-actin | aatcgtgcgtgacatcaaag | ggattccatacccaagaagg |
| Human β-actin | tcaccatggatgatgatatcgc | aatccttctgacccatgcc |
